# 3D multiplexed tissue imaging reconstruction and optimized region-of-interest (ROI) selection through deep learning model of channels embedding

**DOI:** 10.1101/2022.12.09.519807

**Authors:** Erik Burlingame, Luke Ternes, Jia-Ren Lin, Yu-An Chen, Eun Na Kim, Joe W. Gray, Sandro Santagata, Peter K. Sorger, Young Hwan Chang

## Abstract

Tissue-based sampling and diagnosis are defined as the extraction of information from certain limited spaces and its diagnostic significance of a certain object. Pathologists deal with issues related to tumor heterogeneity since analyzing a single sample does not necessarily capture a representative depiction of cancer, and a tissue biopsy usually only presents a small fraction of the tumor. Many multiplex tissue imaging platforms (MTIs) make the assumption that tissue microarrays (TMAs) containing small core samples of 2-dimensional (2D) tissue sections are a good approximation of bulk tumors although tumors are not 2D. However, emerging whole slide imaging (WSI) or 3D tumor atlases that employ MTIs like cyclic immunofluorescence (CyCIF) strongly challenge this assumption. In spite of the additional insight gathered by measuring the tumor microenvironment in WSI or 3D, it can be prohibitively expensive and time-consuming to process tens or hundreds of tissue sections with CyCIF. Even when resources are not limited, the criteria for region-of-interest (ROI) selection in tissues for downstream analysis remain largely qualitative and subjective as stratified sampling requires the knowledge of objects and evaluates their features. Despite the fact TMAs fail to adequately approximate whole tissue features, a theoretical subsampling of tissue exists that can best represent the tumor in the whole slide image. To address these challenges, we propose deep learning approaches to learn multi-modal image translation tasks from two aspects: 1) generative modeling approach to reconstruct 3D CyCIF representation and 2) co-embedding CyCIF image and Hematoxylin and Eosin (H&E) section to learn multi-modal mappings by a cross-domain translation for minimum representative ROI selection. We demonstrate that generative modeling enables a 3D virtual CyCIF reconstruction of a colorectal cancer specimen given a small subset of the imaging data at training time. By co-embedding histology and MTI features, we propose a simple convex optimization for objective ROI selection. We demonstrate the potential application of ROI selection and the efficiency of its performance with respect to cellular heterogeneity.

## INTRODUCTION

Cancers are complex diseases that operate at multiple biological scales—from atom to organism—and the purview of cancer systems biology is to integrate information between scales to derive insight into their mechanisms and therapeutic vulnerabilities. From this holistic perspective, the field has come to appreciate that the spatial context of the tumor microenvironment in intact tissues enables a more granular definition of disease and the design of more personalized and effective therapies^1^. This has been spurred by an increased understanding that solid tumors are complex ecosystems including stromal barriers imposed by tissue architecture^2^ and infiltrating immune cells in the surrounding stroma^3^. This has motivated the National Cancer Institute’s Human Tumor Atlas Network (HTAN) to begin charting 3D tissue atlases which capture the multiscale organizations and interactions of immune, tumor, and stromal cells in their anatomically native states^4^. The HTAN-SARDANA^5^ is one such atlas that aimed to deeply characterize the architecture of a single colorectal cancer (CRC) specimen via histology and a spatial context-preserving multiplexed imaging platform called cyclic immunofluorescence (CyCIF)^6^.

Histology is an essential component of the clinical management of cancer. For around 150 years, pathologists have interrogated thin sections of tissue stained with hematoxylin and eosin (H&E) to determine the morphological correlates of cancer grade, stage, and prognosis. However, this essentially 2D representation of tissue is a relatively poor representation of tissues like the prostate, pancreas, breast, and colon which have highly convoluted 3D ductal structures^5,7–9^. Since 2D whole slide imaging of a 3D specimen might not be representative, 2D analyses using biased down-sampling or the small fields of view afforded by tissue microarrays (TMAs) suffer further due to subsampling issues^5,10^. Moreover, histology alone lacks the molecular specificity to unequivocally determine the identity and function of cells in tissue. In contrast, CyCIF enables the co-labeling of tens of markers in tissue and can broadly characterize the tumor, immune, and stromal compartments. By coupling histology and CyCIF in the same specimen, the HTAN-SARDANA atlas integrates both top-down (pathology-driven) and bottom-up (single-cell phenotype-driven) perspectives of CRC and provides a framework for the charting of 3D atlases for other cancers^5^.

In spite of these advances, 3D multiplexed imaging atlases and 2D whole slide multiplexed imaging with large cohorts both require a tremendous amount of resources and effort to build. For the HTAN-SARDANA atlas, a single CRC specimen was serially sectioned and processed yielding 22 H&E slides interleaved with 25 CyCIF slides, with the CyCIF slides taking days to process due to the cycles of antibody incubation. To build the breast cancer atlas, a single specimen was serially sectioned and processed into 156 slides which were characterized using imaging mass cytometry^8^, which enables simultaneous labeling of 40 antigens with a single incubation step, but has a relatively limited spatial scope (500 μm x 500 μm x 500 μm) compared to CyCIF. To build the pancreas cancer atlas, specimens were serially sectioned and processed into over 1,000 H&E slides, some of which had histological regions of interest labeled through a laborious and subjective manual annotation process^9^. These annotations were used as training data for a deep learning segmentation model which was used to fully reconstruct the semantically-labeled 3D specimen with high accuracy, but this approach is restricted by the limited and predefined annotation classes.

To address this challenge, we extend a virtual staining paradigm into the third dimension by deploying it on the coupled H&E and CyCIF image data from the HTAN-SARDANA atlas of CRC. We have previously demonstrated methods for predicting virtual IF stains based on H&E-stained tissue (SHIFT: Speedy Histological-to-ImmunoFluorescent Translation)^11,12^, wherein we use spatially-registered H&E and immunofluorescence (IF) data and generative deep learning to model the correspondences between these imaging modes and compute near-real time virtual IF stains conditioned on H&E-stained tissue alone. From a biological perspective, these data and approaches allow us to ask which markers in an IF panel have a quantifiable histological signature, what that signature might be, and a means to estimate the distribution of markers in histological images for which such a signature exists. From an application perspective, the approach could be useful for automated compartment labeling in 3D tissues labeled with highly-standardized and low-cost histological stains. We demonstrate that what generative models learn from less than 5% of coupled H&E and CyCIF images is sufficient to generate a virtual 3D CyCIF reconstruction of the whole CRC specimen and that quantitative endpoints derived from real and virtual CyCIF images are highly correlated.

In order to reduce the burden and complexity of multiplex imaging on whole slide images (WSIs), TMAs are often used to sample small sections of the tissue for analysis. Although these TMAs have become a staple of analytics over the past decade, they come with many drawbacks and are prone to substantial bias, often introducing sampling errors and shifts in the expected content which fail to accurately capture the true heterogeneity and spatial distributions found in WSIs^10,13^. In order to overcome this sampling bias, a significantly large number of TMA cores would need to be taken^10,14^, but increasing the size of the randomly sampled TMA cores also shows little to no effect on improving their representativeness^15^. It is necessary to intelligently sample regions for TMAs, but without a method to quantify biological content beforehand, intelligent sampling is estimated from histological appearance alone. If regions of WSIs could be quantitatively described prior to analysis, TMA cores could subsequently be taken based on which regions of the image were most representative of the whole slide.

As a method for virtual TMA selection, we further explore the concept of shared representation between H&E and CyCIF to quantitatively identify representative samples for a region of interest (ROI) selection. Using the principles of SHIFT^11,12^, here we propose a cross-domain autoencoder (XAE) image translation architecture which after training can assign regional descriptors to image tiles that contain the cell type information of CyCIF based solely on the H&E image. By formulating a simple convex optimization problem, these tile-based descriptors can be used to select small regions that are representative of the whole slide image with a minimum number of ROIs. We demonstrate a proof-of-concept study that the XAE architecture is able to adequately represent biological information and that the minimum set of ROIs is more representative of whole slide biology than random sampling or biased manual ROI selection.

## RESULTS

### Preprocessing steps for spatially registered H&E and CyCIF images

Spatially registered H&E and IF images are a requirement for SHIFT model^12^ training and evaluation. To register the H&E and CyCIF data for this task, we begin with sequential registration of the H&E stack beginning from the middle sections and propagating to outer sections (see Methods section, Supplementary Figure 1A and B). We then co-register ROIs of adjacent H&E and CyCIF images (5 μm apart) using their respective nuclear masks for a finer local registration of the adjacent sections.

Before SHIFT model training could begin, we had to account for the section-to-section variability in H&E stain intensity, which helps to ensure a model trained on one H&E section generalizes well to the other sections. Using the training H&E section (middle section as shown in Supplementary Figure 1A) as a reference, we tried several stain normalization methods for outer testing sections^16–18^, and found that the Reinhard method worked best at normalizing stain intensities to the reference by qualitative comparison (Supplementary Figure 1C and D). This result was consistent with a quantitative comparison that found the Reinhard method conferred better generalizability to deep learning models in an analogous digital pathology application^19^.

### Image-to-image translation for 3D virtual CyCIF reconstruction

With spatially registered H&E and CyCIF data, we set out to generate a virtual 3D CyCIF reconstruction in an effort to measure how faithfully we can characterize the full SARDANA dataset with virtual IF staining by learning from only one pair of adjacent H&E and real CyCIF sections. First, the middle pair of H&E and CyCIF sections were selected for training SHIFT models under the assumption that they are a good representation of the tissue on either side of the sample block. This assumption is supported by the initial HTAN-SARDANA study^5^, where the authors conclude that 2D whole slide imaging of a 3D specimen does not, in general, suffer from the subsampling issue associated with TMAs or small fields of view.

We then decompose the WSIs into thousands of pairs of matching H&E and CyCIF image tiles and use those to train a generative adversarial network (GAN) to synthesize virtual CyCIF tiles conditioned on H&E tiles^12^. Briefly, the generator network of the model is responsible for synthesizing virtual CyCIF images conditioned on H&E images, and the discriminator network is responsible for quality assurance of the virtual CyCIF images synthesized by the generator as shown in Figure 1A. Once trained on the middle sections, the model can then be tested by feeding it tiles from the held-out H&E sections to generate virtual CyCIF images for comparison with the real CyCIF images. Importantly, a virtual CyCIF image is conditioned on H&E section, and there is natural variation between it and its adjacent real CyCIF section 5 μm away, which complicates pixel-wise evaluation of model accuracy.

**Figure 1:**
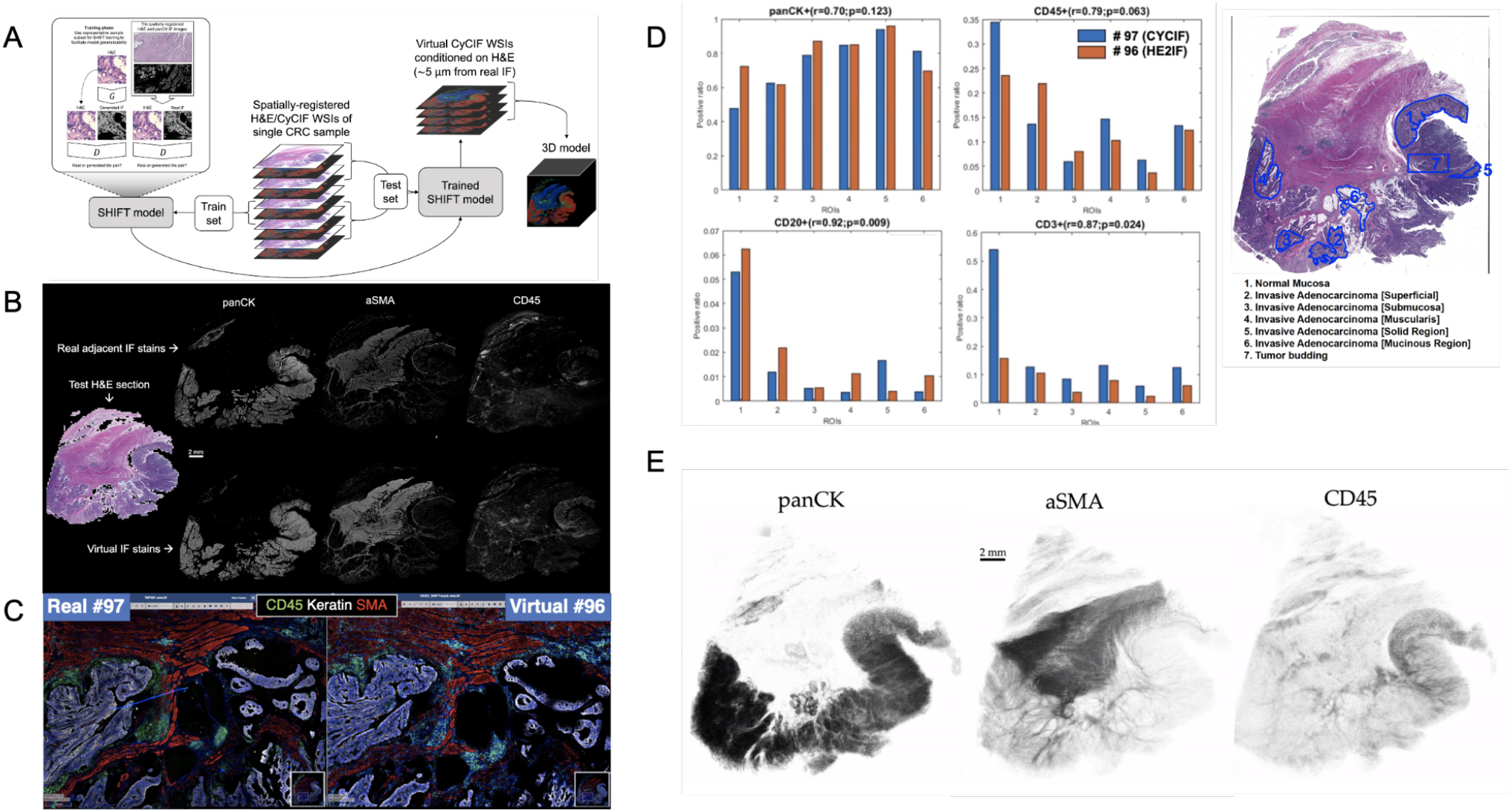
Overview of Image-to-Image translation for 3D virtual CyCIF reconstruction of SARDANA and WSI virtual staining result: (A) Extending SHIFT to 3D using adjacent spatially-registered H&E/CyCIF WSIs from a single CRC sample. (B) WSI virtual staining result. Models trained to predict single-channel CyCIF images conditioned on the H&E/CyCIF training sections were applied to H&E test section 096 to generate virtual stain WSIs for the markers panCK, αSMA, and CD45. The input H&E test section is shown at left, and the real and virtual CyCIF WSIs are shown in the rows above and below, respectively, for ease in comparison. (C) qualitative comparison of real and virtual staining for the markers panCK, αSMA and CD45 in the selected region. (D) Quantitative comparison of ROI cell composition correlation between real. For each of the ROIs, the positive ratio of cells for each of panCK, CD45, CD20, and CD3 are calculated using the same workflow and displayed for either real or virtual CyCIF WSIs. Pearson’s correlations and p-values describing the association between positive ratios derived from real and virtual CyCIF WSIs for each marker are indicated above each bar plot. (E) 3D virtual stain volumes conditioned on held-out H&E test sections visualized by 3D Slicer^39^.

We trained individual SHIFT models to predict single CyCIF channels conditioned on H&E inputs from the central H&E/CyCIF training sections 053/054 (Figure 1A). Representative test results from the application of trained SHIFT models on H&E/CyCIF test sections 096/097 (far from the middle section, i.e., training section) are shown in Figure 1B and 1C. These qualitative results indicated that the SHIFT models were fitting well with the training sections, and the representations learned were useful for an extension to held-out test sections.

The virtual CyCIF images generated by SHIFT models are conditioned on H&E sections which are 5 μm adjacent to the real CyCIF sections, so the cellular contents are slightly different between sections and images. Recognizing that this would hamper pixel-wise comparisons between the real and virtual CyCIF images^11,12^, we estimated an upper bound on SHIFT performance by measuring the concordance between nuclear content from the adjacent sections of the H&E/CyCIF test sections 096/097 (Supplementary Figure 2).

The test sections were first subdivided into 135 non-overlapping ROIs and each ROI was locally registered to improve the alignment of H&E and CyCIF image content, then we measured the Dice coefficient of nuclear masks derived from the H&E and DAPI images from each ROI (Supplementary Figure 2A). We used the Dice coefficient for each ROI as a compensation factor when evaluating the quality of the virtual stains for each ROI by dividing raw quality scores by the Dice coefficients corresponding to each ROI. Virtual CyCIF image quality was evaluated using structural similarity (SSIM), which is established as a metric for assessing virtual stain quality^12,20,21^. The median compensated SSIM for virtual stains ranged from 0.36 for CD20 up to 0.89 for αSMA. This result suggested that there was significant room for improvement for some SHIFT models, but we hypothesized that the virtual images might still be useful in the hands of a CyCIF domain expert since SSIM is sensitive to slight differences in image contrast which may not significantly affect downstream processing and interpretation^12^.

To test this, we quantified the positive cell ratio for multiple markers in each of the pathologist-annotated 6 ROIs in H&E test section 096 using either real or virtual CyCIF images (Figure 1D), which assesses how such an endpoint might be impacted when using virtual images which may or may not be of high quality with respect to SSIM (Supplementary Figure 2). In spite of the adjacency complication explained above, there was a substantial correlation between positive cell ratios using real and virtual CyCIF images, suggesting that virtual images could be used in place of real without significantly affecting some downstream endpoints. Having established the fitness of the SHIFT models, we performed a full virtual 3D reconstruction of the CyCIF images by passing all held-out H&E test sections to the SHIFT models trained on the H&E/CyCIF training sections (Figure 1E).

We also tested ablation study to assess the value added by the discriminator network of the GAN by training models without it, leaving the generator network to learn the virtual panCK stain alone (Supplementary Figure 3). We found that while the generator-only virtual panCK stain has good localization, it lacks the naturalistic texture of the real and GAN-generated virtual stains, which highlights the compromise of a more efficient and portable generator-only model.

### Shared latent representation via embedding of CyCIF images on H&E image

3D Virtual staining is enabled through the rich latent representations that generative models are capable of learning from paired H&E and CyCIF image data. We hypothesized that these latent representations could be useful for the related and unsolved problem of objective ROI selection. If ROI selection for targeted CyCIF staining was to be possible using only H&E for prediction, it would be necessary for the H&E images to contain relevant biological information equivalent to that of CyCIF.

To test this hypothesis, we created tile-based image descriptors from H&E using a standard Variational Autoencoder (VAE)^22^ and compared them to cell type composition vectors (7 cell types) created from CyCIF imaging data for the same tiles. In order to evaluate the overlap and exclusivity of each modality’s information, we used canonical correlation analysis (CCA)^23^ using two components. The two modalities quantitatively show canonical correlations (0.91 and 0.88 for each component respectively), and qualitatively show a high level of overlap when the two components are plotted on top of one another (Figure 2A). Motivated by this example, and building upon previous works in cross-domain data translation^24,25^, we built a cross-domain autoencoder (XAE) architecture that learns to co-embed H&E and CyCIF representations of the same tissue into the shared latent representation (Figure 2B). To test a minimum working example of our XAE architecture, we performed a simple ablation experiment with the CyCIF encoder of the model removed. For this experiment, the model was tasked with H&E reconstruction and H&E-to-(DAPI and panCK) translation. To assess the goodness of fit, the model was trained to convergence and evaluated on a training batch. Visual inspection of model outputs indicated that the model was functioning as intended (Figure 2C). In our original design, the XAE included skip connections that connected across the U-Net generator blocks, but we discovered that the models did not learn useful latent representations of images, a direct effect of the absence of loss function gradient flow through the interior layers of the models enabled by skip connections. We removed the skip connections in subsequent experiments and found that these models exhibit good convergence properties and have appreciable loss function gradient flow through the model interior (not shown).

**Figure 2.**
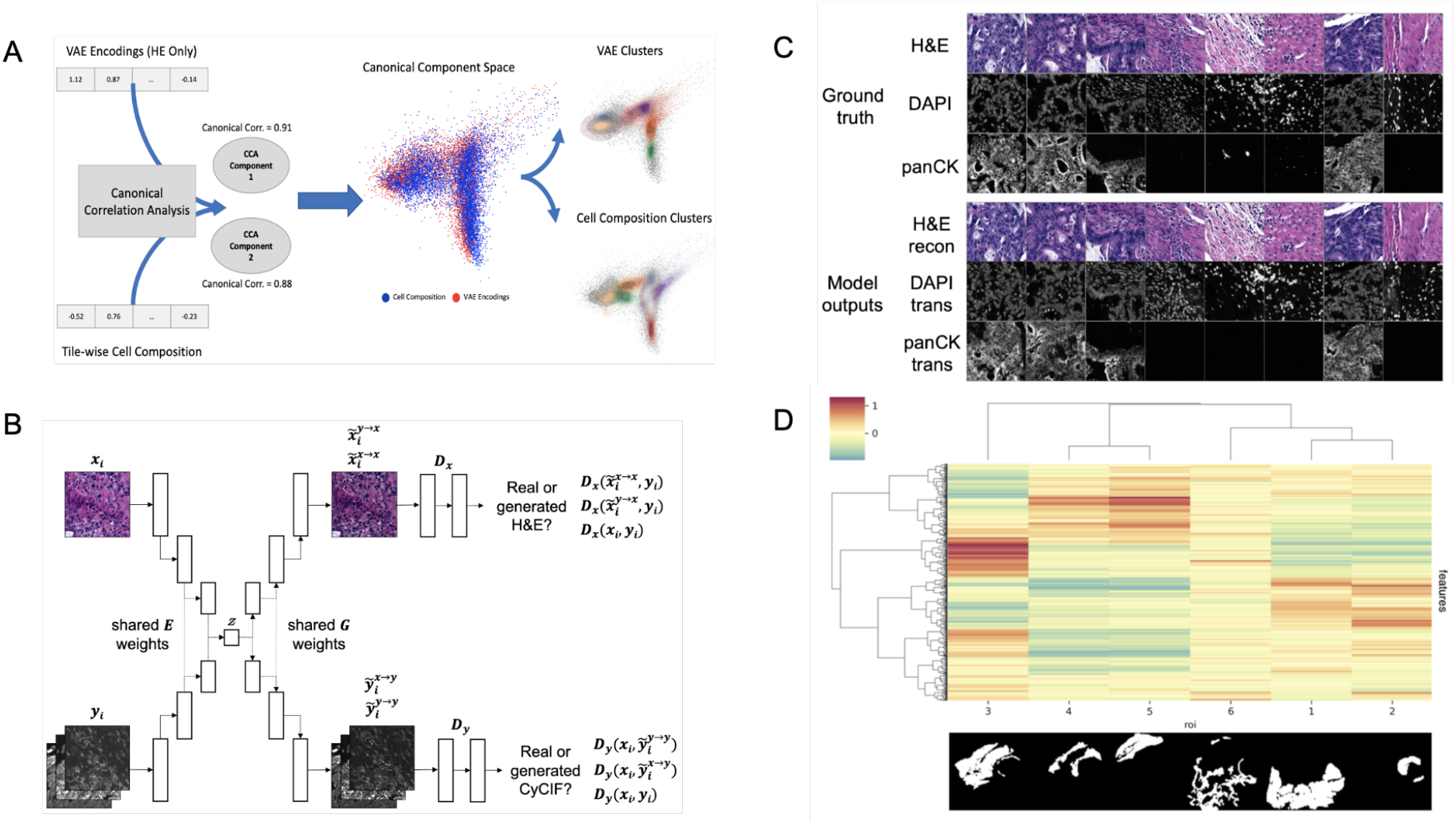
(A) VAE encodings of HE and CyCIF cell type composition (7 cell types) show high canonical correlation and a large overlap between data and cluster embeddings. (B) XAE architecture. The model has two input heads, one for H&E encoder inputs (xi) and another for CyCIF encoder inputs (yi), both of which encode into a shared latent space (z). The model also has two output heads, one for H&E decoder outputs and another for CyCIF decoder outputs. Full XAE model architecture is described in Table 2. (C) Ground truth tiles representing a single training batch. Trained XAE model results for the tasks of H&E-to-H&E reconstruction and H&E-to-CyCIF translation using the ground truth training. (D) XAE latent feature clustering and corresponding pathologist annotation where the inset image indicates the binary mask corresponding to each ROI with respect to the layout of the H&E test section. Features were z-scored, then tiles were mean-aggregated based on their ROI, and features were hierarchically clustered. The ROI label keys are 1: tumor adenocarcinoma (n = 2501 tiles); 2: normal mucosa (n = 362 tiles); 3: proper muscle (n = 1576 tiles); 4: submucosa (n = 473 tiles); 5: subserosa, loose connective tissue (n = 782 tiles); and 6: fibrosis, inflammation, lymphoid aggregate (n = 1048 tiles). The color scale corresponds to the mean of z-scored feature values for each ROI.

Having confirmed that the trained XAE had fit its training distribution (Figure 2C), we next wanted to assess the representativeness and interpretability of the latent feature space that it learned with respect to pathologically interesting regions of the sample. To do this, we used the H&E encoder of the trained XAE to encode tiles from H&E test section 096 into 512-dimension feature representations and assessed how the features were distributed over tiles drawn from each of several pathologist-defined ROIs in the test section. The 6742 nonoverlapping tiles from H&E test section 096 which had at least one pixel of pathologist annotation were each encoded into 512-dimension latent feature maps. We found that many of the learned image features were associated with pathologically-distinct regions of the sample (Figure 2D).

In order to evaluate how well deep learning can capture and represent unseen complex information using H&E images alone, the VAE model trained on H&E images alone and XAE features were compared to cell types defined by CyCIF expressions and pathologist tissue annotations. Clustering tiles within the whole slide image based on cell type composition using K-means resulted in 7 clusters, and the pathologist annotated 6 key tissue types to be used as ground truth (Figure 3A). Ground truth tile labels were compared against one another to create a baseline for evaluation. When annotations were used to predict cell type, there was a baseline performance of 57.1% cluster purity and 0.44 normalized mutual information (NMI). Conversely, when cell type was used to predict annotations, there was a baseline performance of 66.8% cluster purity and 0.44 NMI (Figure 3B). In all metrics, XAE outperformed VAE predictions, achieving a 56.1% cluster purity and 0.35 NMI against cell type, and 70.2% cluster purity 0.38 NMI against pathologist annotation (Figure 3B). It is also notable that on the metric of cluster purity against annotations, the XAE outperformed the baseline metric; this indicates that the XAE is better at predicting histologic tissue type than even cell type compositions.

**Figure 3:**
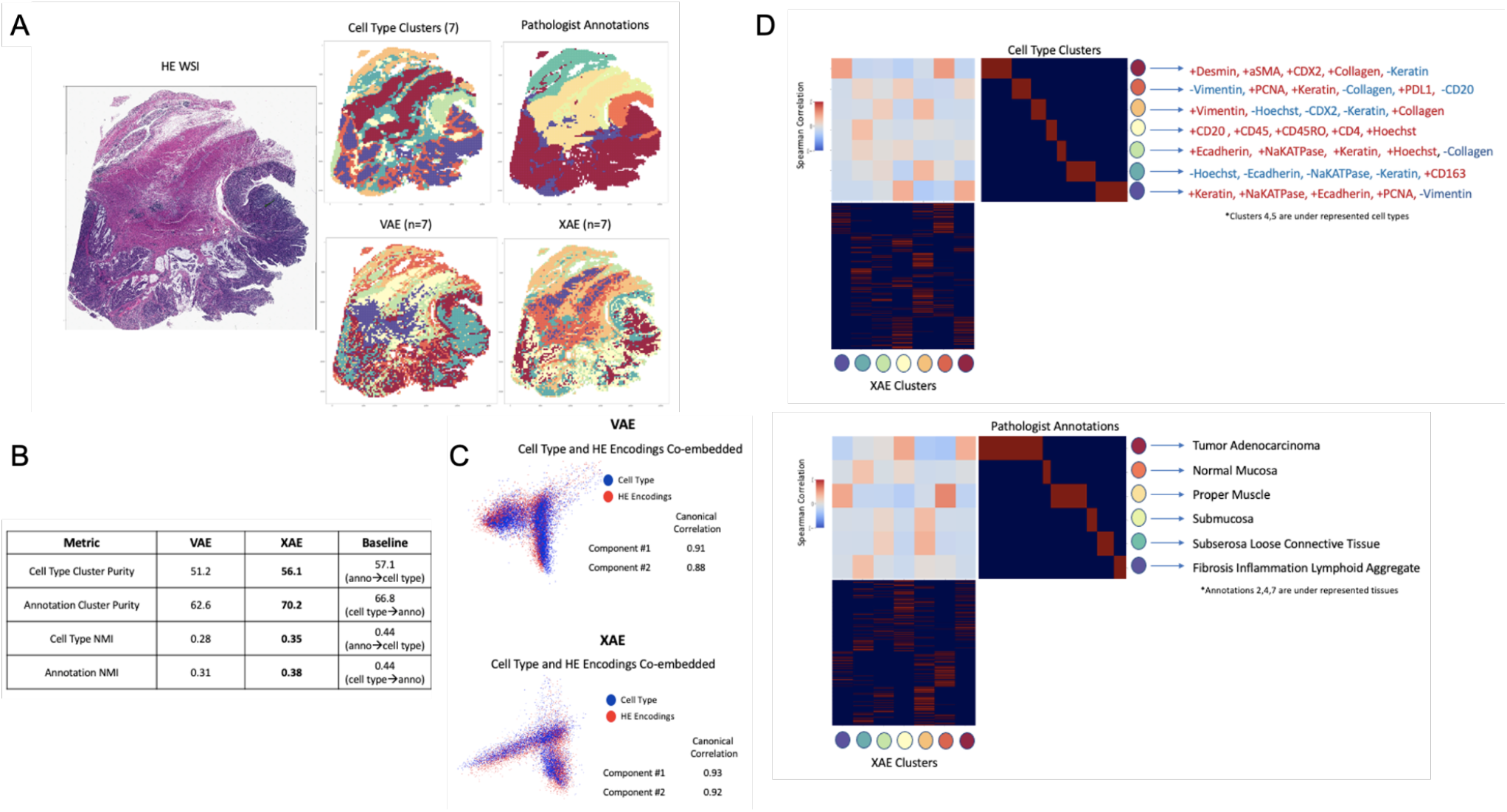
Deep learning architectures recapitulate unseen complex information using H&E. (A) Images colored by tile labels for cell type, pathologist annotation, assigned cluster from VAE using H&E input, and assigned cluster from XAE using H&E input. (B) Quantitative evaluation of VAE and XAE at recapitulating biological labels, measured using cluster purity and NMI and compared to baseline of agreement between biological labels. (C) Canonical correlation analysis between cell type composition vector and H&E encodings for both VAE and XAE, quantitatively measured by component correlation and qualitatively by label overlap in embedding space. (D) Cluster-wise correlation matrix for XAE against both cell type and pathologist annotations to determine which biological features are adequately captured. Defining CyCIF expressions provided based on inter/intra-cluster variability.

Analysis of complex information, deeper than large-scale clustering, was conducted using canonical correlations between the model embedding space and the tile-wise CyCIF expressions. Visually both VAE and XAE show a good overlap between cell type embeddings from CyCIF and model embeddings produced from H&E images (Figure 3C); the XAE, however, achieves higher canonical correlations (0.93 and 0.92 compared to 0.91 and 0.88 for VAE). To confirm that we were extracting relevant and rare cell types with the representation models, we computed the Spearman correlation between every predicted cluster and the ground truth cluster (Figure 3D). From this, we can see that XAE has consistently high magnitudes of correlation and that a reasonable correlation exists for every ground truth cluster except for cell type clusters 4 and 5 which are underrepresented populations. Furthermore, the cell types that the XAE is able to capture are largely explained by changes in Na-K ATPase, E-Cadherin, and PCNA, which were shown to be important indicators for cell phenotypes in prior research on this tissue^5^.

It is shown by numerous metrics that the XAE model outperforms the VAE in capturing detailed information from H&E images alone, which are able to adequately recapitulate information from CyCIF expression data and pathologist annotations that are unseen during test time. Because the XAE encodings are able to adequately recapitulate the information in CyCIF from H&E, we can use them for proxy analyses such as selecting representative regions of the WSI for further analysis.

### Co-embedding H&E and CyCIF representations improve ROI selection

Currently, ROI selection within H&E WSIs is done either randomly, which is inaccurate and is likely to select an area that doesn’t represent the WSI, or with manual selection of ROI, which is biased, subjective, and has been shown to miss whole tissue patterns^5^. Using the XAE embeddings, which capture the complex cell type and annotation information using H&E, we develop an optimization-based approach to select a minimum set of ROIs that are more representative than random sampling while being repeatable and biologically driven. To evaluate ROI selection performance, we use three metrics: mean squared error (MSE) between the cell type composition of selected ROIs and WSI; Jensen-Shannon Divergence (JSD) between the cell type composition vectors of selected ROIs and WSI; and mean entropy of the selected ROIs’ cell type compositions. Since MSE and JSD both have disadvantages, the use of both for evaluating composition is beneficial. MSE is highly prone to outliers and abnormal data, amplifying errors of single erroneous samples, and JSD cannot operate with terms that are zero (ignoring them from the operation), and therefore underestimates error in samples with empty classes. Three different methods for ROI selection were tested: random sampling, convex optimization minimizing *l*_1_-norm of cell type composition, and convex optimization minimizing the norm of cell type composition with maximizing entropy to select ROIs with more heterogeneous cell composition.

When regions are randomly sampled, we observe that the cell type compositions struggle to converge to the whole slide cell type composition, taking upwards of 20-30 ROIs (each of which comprises between ~0.15% and ~0.80% of WSI area individually) before reaching a reasonable representation (Figure 4 top row). Using a simple composition-based optimization, selected ROIs drastically decrease the number of ROIs necessary to around 7. This number of ROI is equivalent to the number of cell type clusters we were optimizing for and further investigation shows that the algorithm was selecting primarily homogeneous regions that reconstruct the whole slide composition. This is validated by looking at the mean entropy of ROIs for the base convex optimization method, which consistently shows low to middling ROI entropy values, especially in the 1000-pixel size data (Figure 4 middle row).

**Figure 4:**
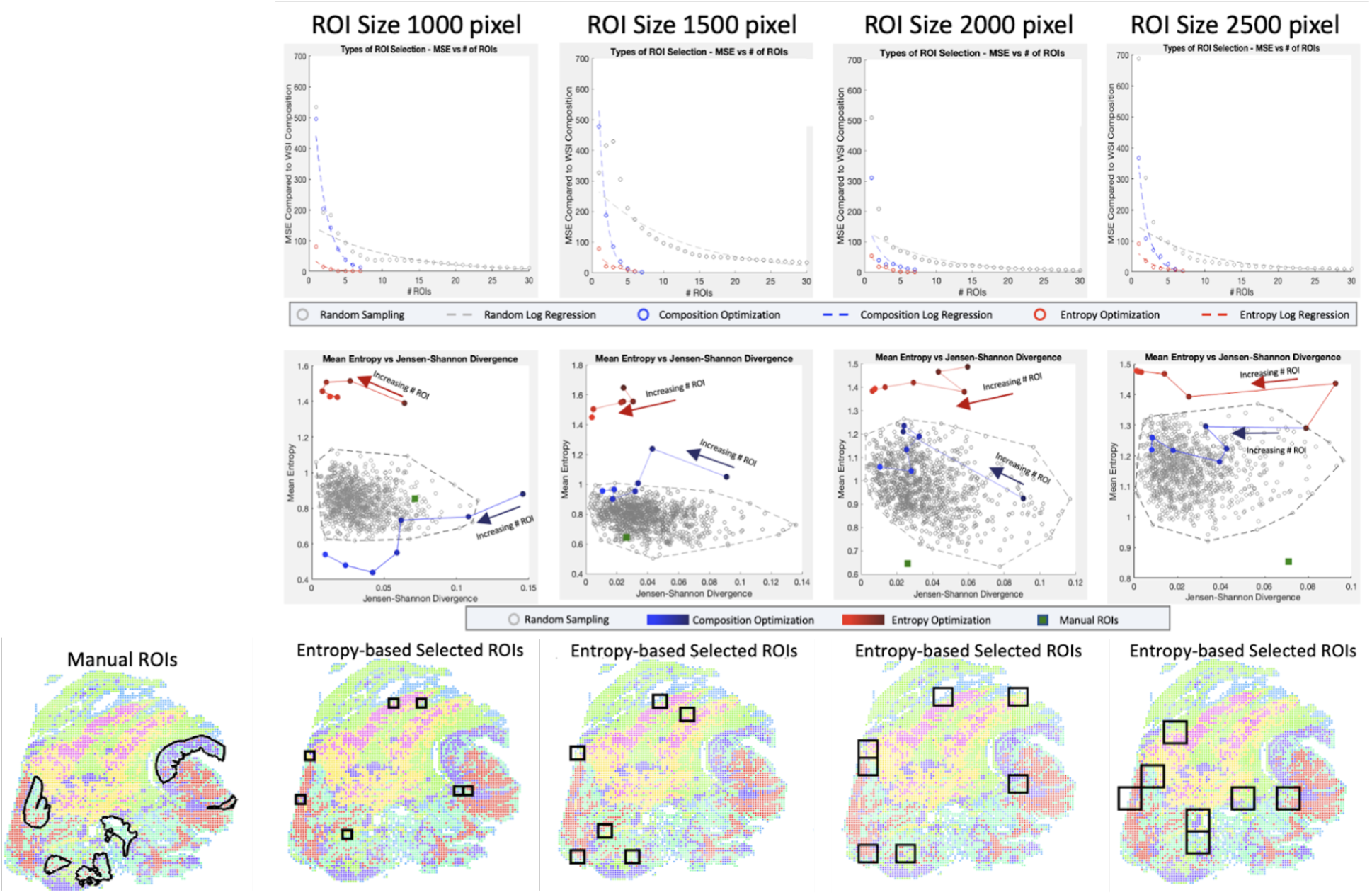
Optimization of ROI Selection. For four ROI sizes (1000×1000, 1500×1500, 2000×2000, 2500×2500 pixels) and three sampling techniques (random sampling, convex optimization using cell type composition, convex optimization using cell type composition, and regional entropy), we calculate the optimal selection of ROI. (Top row) By calculating the MSE for a range of ROI, we can evaluate each technique’s rate and quality of convergence. (Middle row) Selections of representative ROIs are evaluated based on two metrics (Entropy for tissue heterogeneity and Jensen-Shannon Divergence for composition similarity). Random sets of 7 ROIs are generated 1000 times to portray the baseline pattern. Selections from linear and convex optimizations are plotted with increasing numbers of ROIs to show the change in performance. The performance of the manually selected ROIs is also shown to emphasize the bias in targeted sampling. (Bottom row) The optimal ROIs are shown for convex entropy optimization at each size of ROI. Image colors portray the XAE labeled cell types.

To select a more heterogeneous region, entropy is considered in the convex optimization and we observe convergence much earlier at 3-4 representative ROIs. Unlike the simple optimization considering cell composition only, however, the ROIs selected are not homogenous and include much more biologically interesting regions with diverse cell populations. This is confirmed by entropy values considerably higher than the randomly sampled population. When looking at the full range of clusters, both optimization-based approaches are substantially better than even manual ROI selection which is extremely biased, scoring poorly on both composition metrics and heterogeneity metrics.

To account for this, we narrowed the range of clusters being optimized for in the ROI selection to only consider tumor and immune cell populations (Supplementary Figure 4). Even in this restricted cluster set, manual annotation does not perform better than convex optimization using entropy and is less representative of the WSI’s tumor and immune cell type composition. This shows that the improvements made over manual selection are not solely due to the cell type bias of pathologists selecting interesting regions; it is also the fact that the ROI selection based on the convex optimization method can find the most representative regions which can be a difficult task for an annotator who cannot see cell type.

## DISCUSSION

Tumors are not 2D, but many of the imaging characterization platforms in both research and clinical practice make the assumption that TMAs containing small core samples of essentially 2D tissue sections are a reasonable approximation of bulk tumors. However, emerging 3D tumor atlases strongly challenge this assumption^5,26,27^. In spite of the additional insight gathered by measuring the tumor microenvironment in 3D, it can be prohibitively expensive and time-consuming to process tens or hundreds of tissue sections with CyCIF. Even when resources or time are not limiting, the criteria for ROI selection in tissues for downstream analysis remain largely qualitative and subjective.

In the current study, we extend the virtual staining paradigm to a 3D CRC atlas^5^ and demonstrate a proof-of-concept that generative models can learn from a minimal subset of the atlas to reconstruct the remaining sections of the CyCIF portion of the 3D atlas and recapitulate the quantitative endpoints derived using the real CyCIF data. Quantitative comparisons of real and virtual CyCIF stains exposed the challenge of using adjacent sections to train models, where image contents are subtly but appreciably different between sections at single-cell resolution. This challenge could be overcome in future studies by staining each tissue section first with CyCIF, then terminally with H&E^12,28^. That being said, this study and those like it take for granted that histology workflows are inherently destructive since serial sectioning and processing of tissue can preclude tissue from being used in other assays. Alternatively, a non-destructive 3D microscopy approach using tissue clearing and light-sheet microscopy could be deployed, which would also preserve tissues for other assays^7^. However, the slow diffusion rate of antibodies in whole tissues limits the deep multiplexing potential of the CyCIF platform in this non-destructive approach, but the use of small molecule dyes and affinity agents could help to overcome this challenge to 3D virtual staining applications^29^.

We also implement and evaluate a novel deep learning model which integrates paired H&E and CyCIF data into a shared representation and demonstrate that the model can be used as a quantitative and objective guide for ROI selection, with the integrated H&E/CyCIF representations being more informative than H&E representations alone. The limitation of this approach is that the XAE model must be trained using paired H&E-CyCIF data prior to being used for prediction and quantification but we can also reduce the required CyCIF panel^30^. A further limitation is that the ROI selection can only be optimized with respect to quantifiable measures such as heterogeneity and composition.

Although image representations can accurately describe biological features, they cannot convey what may or may not be biologically interesting to researchers or clinicians. Although cell type composition and entropy were used as metrics of biological relevance in this setting, it is likely that other experiments would have different priorities. Some examples of this might include: weighting cell type clusters by the level of interest; weighting entropy negatively if homogeneous regions are desired; weighting some other extracted scores such as colocalization of two cell types of interest. The method of optimization is versatile and amenable to many different functions. The key takeaway is that this pipeline allows for intelligent representation from H&E images, which enables a plethora of subsequent analyses on this representation space with other multiplexed imaging platforms such as multiplexed ion beam imaging (MIBI)^31^, imaging mass cytometry (IMC)^32^, or NanoString GeoMX^33^ as only a few ROIs could be selected and analyzed using these platforms.

## METHODS

### 3D registration of paired H&E and CyCIF

Because images are taken on serial sections, images throughout the 3D stack of tissue and between H&E and CyCIF require registration in order to be properly analyzed. To register all the H&E together, we used the centermost slide as the baseline target for registration (i.e., reference). Registration transforms were calculated between each layer in the stack, then were applied sequentially to all slides, moving from one to the next until all slides were registered to the same coordinates as the central slide (Supplementary Figure 1A). The central slide was chosen as the reference because it would maximize similarity to the tissue morphologies at the far ends of the tissue stack.

For training and testing H&E to CyCIF training, it was necessary to have high-quality single-cell level registration of adjacent H&E and CyCIF images. Due to whole slide structural changes that biologically occur in the 5 μm space between sections, it was not possible to adequately register whole slide images this accurately without using non-rigid transformations, which resulted in imaging artifacts that skewed analysis. To get the best registration possible with the least amount of artifacts, we performed fine-tuned CyCIF registration on smaller ROIs covering the entire tissue. Within a single ROI, a rigid transformation can accurately register the tissue without having conflicting transforms from regions located in distant areas of the whole slide. The registration transform for this step was calculated using a binarized DAPI image and a binarized H&E image after deconvolution of the hematoxylin stain to align the nuclei for the two images^34^.

### H&E and CyCIF image intensity normalization

To minimize the influence of technical variability on stain color between H&E sections, we experimented with the application of several stain normalization methods to the H&E WSIs^16–18^ using the Python package stain tools (https://github.com/Peter554/StainTools). To identify and mask out background regions of each WSI (white regions of a slide without tissue), WSIs were each cropped into non-overlapping 256×256-pixel tiles and tiles containing greater than 70% area of pixels with 8-bit intensity greater than (210, 210, 210) were excluded from subsequent normalization steps. To help identify and mask out background pixels in the remaining tiles before model fitting and normalization, the foreground tiles from each H&E WSI were independently standardized such that 5% of all pixels were luminosity saturated. For all normalization methods, we used the H&E WSI from section 054 as the stain reference to which the stain intensity distributions of all other H&E WSIs would fit. After normalizing the foreground tiles of each non-reference WSI to fit the reference stain distribution, tiles were restitched to form cohesive WSIs. On the basis of visual inspection (Supplementary Figure 1B and 1C), we opted to use the Reinhard normalization method, which has also been shown to maximize deep learning model performance on digital pathology applications^19^. To control for variations in raw contrast between CyCIF WSIs, we rescaled the intensities of CyCIF WSIs to have a min-max range fit to the 70th-99.99th intensity percentiles of the input WSIs.

### SHIFT models

SHIFT models were built using Pytorch as previously described^12^. Model architectures are described in Table 1. Models were trained to predict single channel images corresponding to one of the CyCIF stains from input H&E tiles from section 054, e.g. H&E→CD45 or H&E→CD31. Paired H&E and CyCIF image tiles from section 054 were split into 80% training (8134 tiles) and 20% validation (2034 tiles) sets and each model was trained with a batch size of 4 and learning rate of 0.0002 for 100 epochs. Best models were selected based on the lowest validation loss at each epoch end and were then used for downstream application to held-out H&E WSIs.

**Table 1:**
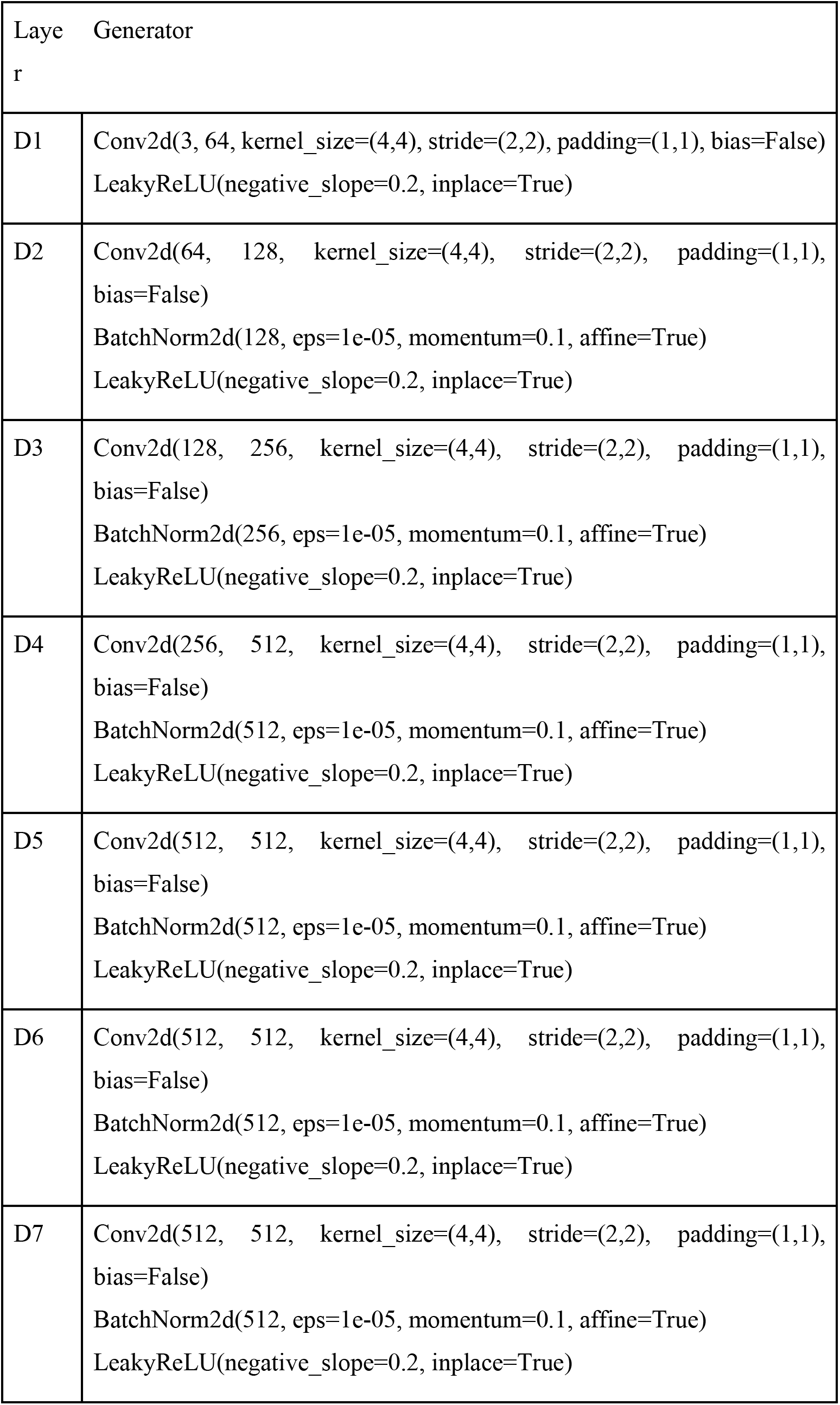
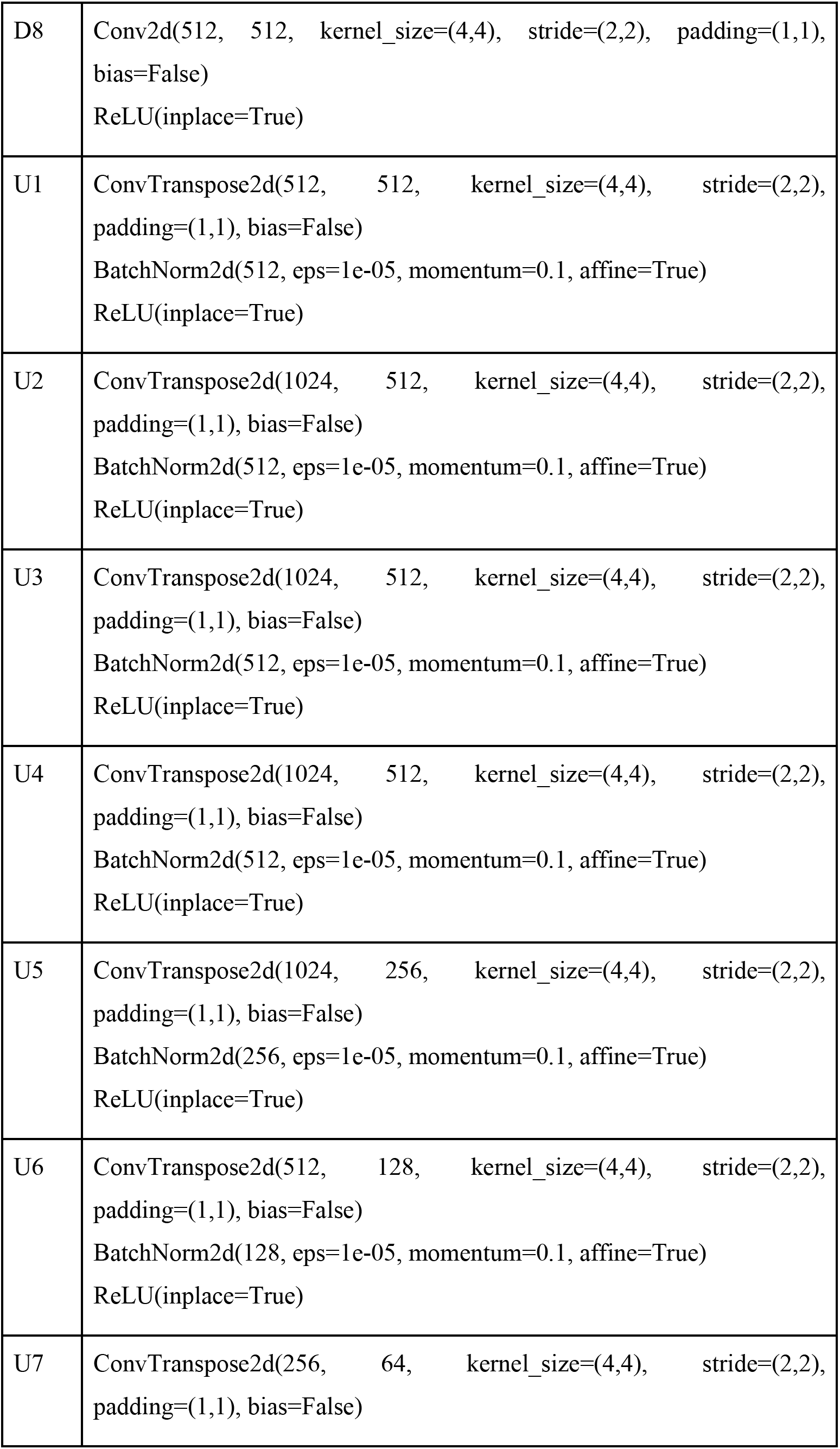
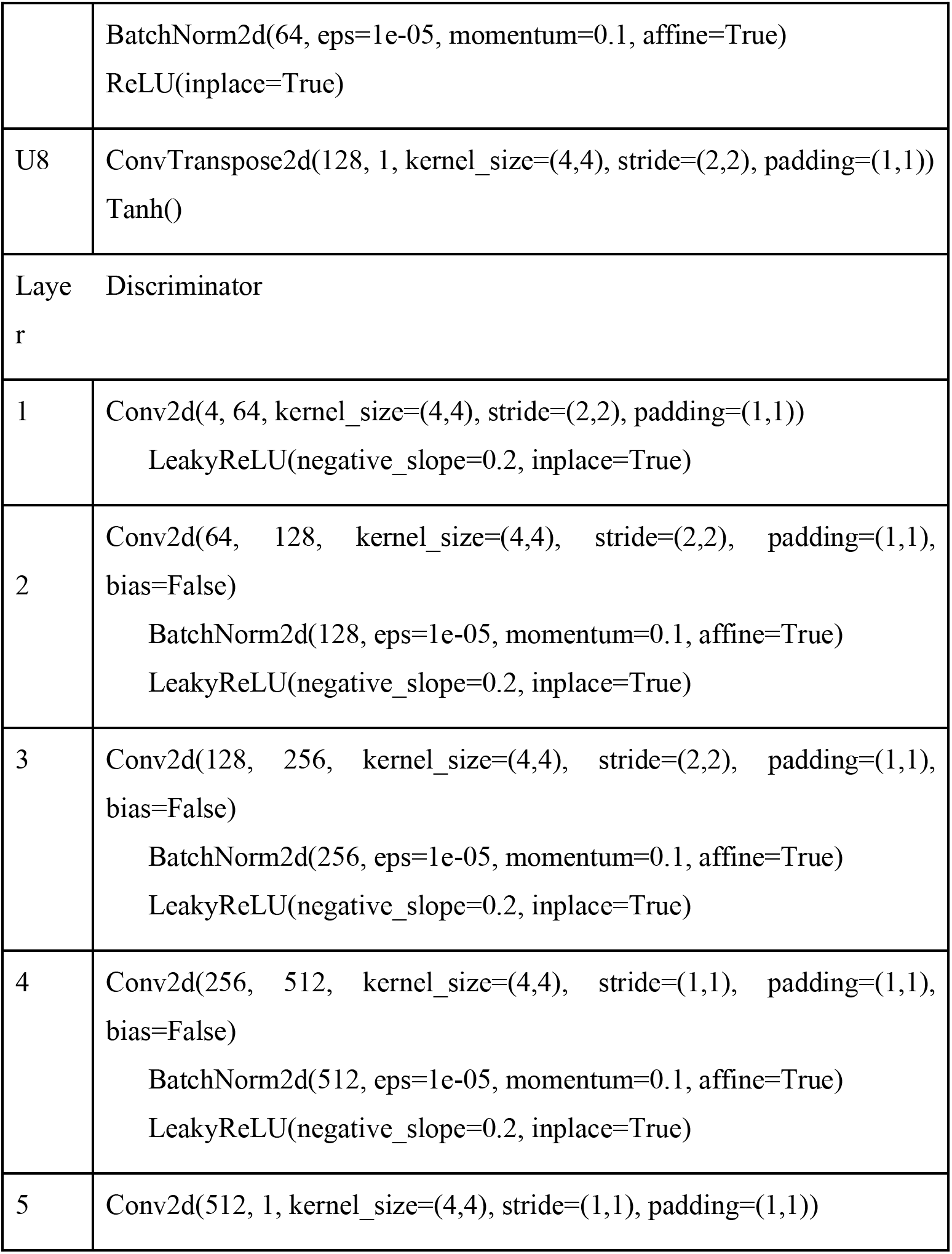
architecture of SHIFT models.

| Layer | Generator |
| --- | --- |
| D1 | Conv2d(3, 64, kernel_size=(4,4), stride=(2,2), padding=(1,1), bias=False)<br>LeakyReLU(negative_slope=0.2, inplace=True) |
| D2 | Conv2d(64, 128, kernel_size=(4,4), stride=(2,2), padding=(1,1), bias=False)<br>BatchNorm2d(128, eps=1e-05, momentum=0.1, affine=True)<br>LeakyReLU(negative_slope=0.2, inplace=True) |
| D3 | Conv2d(128, 256, kernel_size=(4,4), stride=(2,2), padding=(1,1), bias=False)<br>BatchNorm2d(256, eps=1e-05, momentum=0.1, affine=True)<br>LeakyReLU(negative_slope=0.2, inplace=True) |
| D4 | Conv2d(256, 512, kernel_size=(4,4), stride=(2,2), padding=(1,1), bias=False)<br>BatchNorm2d(512, eps=1e-05, momentum=0.1, affine=True)<br>LeakyReLU(negative_slope=0.2, inplace=True) |
| D5 | Conv2d(512, 512, kernel_size=(4,4), stride=(2,2), padding=(1,1), bias=False)<br>BatchNorm2d(512, eps=1e-05, momentum=0.1, affine=True)<br>LeakyReLU(negative_slope=0.2, inplace=True) |
| D6 | Conv2d(512, 512, kernel_size=(4,4), stride=(2,2), padding=(1,1), bias=False)<br>BatchNorm2d(512, eps=1e-05, momentum=0.1, affine=True)<br>LeakyReLU(negative_slope=0.2, inplace=True) |
| D7 | Conv2d(512, 512, kernel_size=(4,4), stride=(2,2), padding=(1,1), bias=False)<br>BatchNorm2d(512, eps=1e-05, momentum=0.1, affine=True)<br>LeakyReLU(negative_slope=0.2, inplace=True) |
| D8 | Conv2d(512, 512, kernel_size=(4,4), stride=(2,2), padding=(1,1), bias=False)<br>ReLU(inplace=True) |
| U1 | ConvTranspose2d(512, 512, kernel_size=(4,4), stride=(2,2), padding=(1,1), bias=False)<br>BatchNorm2d(512, eps=1e-05, momentum=0.1, affine=True)<br>ReLU(inplace=True) |
| U2 | ConvTranspose2d(1024, 512, kernel_size=(4,4), stride=(2,2), padding=(1,1), bias=False)<br>BatchNorm2d(512, eps=1e-05, momentum=0.1, affine=True)<br>ReLU(inplace=True) |
| U3 | ConvTranspose2d(1024, 512, kernel_size=(4,4), stride=(2,2), padding=(1,1), bias=False)<br>BatchNorm2d(512, eps=1e-05, momentum=0.1, affine=True)<br>ReLU(inplace=True) |
| U4 | ConvTranspose2d(1024, 512, kernel_size=(4,4), stride=(2,2), padding=(1,1), bias=False)<br>BatchNorm2d(512, eps=1e-05, momentum=0.1, affine=True)<br>ReLU(inplace=True) |
| U5 | ConvTranspose2d(1024, 256, kernel_size=(4,4), stride=(2,2), padding=(1,1), bias=False)<br>BatchNorm2d(256, eps=1e-05, momentum=0.1, affine=True)<br>ReLU(inplace=True) |
| U6 | ConvTranspose2d(512, 128, kernel_size=(4,4), stride=(2,2), padding=(1,1), bias=False)<br>BatchNorm2d(128, eps=1e-05, momentum=0.1, affine=True)<br>ReLU(inplace=True) |
| U7 | ConvTranspose2d(256, 64, kernel_size=(4,4), stride=(2,2), padding=(1,1), bias=False) |
|  | BatchNorm2d(64, eps=1e-05, momentum=0.1, affine=True)<br>ReLU(inplace=True) |
| U8 | ConvTranspose2d(128, 1, kernel_size=(4,4), stride=(2,2), padding=(1,1))<br>Tanh() |
| Layer Discriminator |  |
| 1 | Conv2d(4, 64, kernel_size=(4,4), stride=(2,2), padding=(1,1))<br>LeakyReLU(negative_slope=0.2, inplace=True) |
| 2 | Conv2d(64, 128, kernel_size=(4,4), stride=(2,2), padding=(1,1),<br>bias=False)<br>BatchNorm2d(128, eps=1e-05, momentum=0.1, affine=True)<br>LeakyReLU(negative_slope=0.2, inplace=True) |
| 3 | Conv2d(128, 256, kernel_size=(4,4), stride=(2,2), padding=(1,1),<br>bias=False)<br>BatchNorm2d(256, eps=1e-05, momentum=0.1, affine=True)<br>LeakyReLU(negative_slope=0.2, inplace=True) |
| 4 | Conv2d(256, 512, kernel_size=(4,4), stride=(1,1), padding=(1,1),<br>bias=False)<br>BatchNorm2d(512, eps=1e-05, momentum=0.1, affine=True)<br>LeakyReLU(negative_slope=0.2, inplace=True) |
| 5 | Conv2d(512, 1, kernel_size=(4,4), stride=(1,1), padding=(1,1)) |

### Measuring concordance between nuclei overlap in adjacent sections

Estimation of the upper bound on SHIFT performance was done by measuring concordance between overlapping nuclei in adjacent sections for locally-registered ROIs from H&E/CyCIF test sections. For H&E ROIs, we deconvolve the hematoxylin stain to extract nuclear content intensity^35^, then segment the intensity to derive binary nuclear masks using Cellpose^36^. For CyCIF ROIs, we use Cellpose to segment DAPI intensity to derive binary nuclear masks. The Dice coefficients describing the overlap of nuclear masks from ROIs of adjacent sections were used as compensation factors for evaluating virtual stains. The Dice-compensated SSIM values are calculated by taking the SSIM (using an 11-pixel sliding window) of the virtual CyCIF ROI with respect to the real CyCIF ROI and dividing it by the Dice coefficient of nuclear overlap between the hematoxylin and DAPI nuclear masks from sections 096/097 for that ROI.

### XAE models

XAE models were built using Pytorch. Model architectures are described in Table 2. The XAE architecture used here is an adaptation of the UNIT architecture^24^ and the imaging-to-omics XAE architecture^25^. XAE models have two input encoders (Figure 2B), one accepting H&E image tiles (batch size × 3 × 256 × 256), and the other accepting the corresponding paired CyCIF images (batch size × N CyCIF channels × 256 × 256). Both encoders compress their inputs into a shared latent space z. From z, image representations can be upscaled by either H&E or CyCIF decoders. Hence, there are four forward paths through the model: (1) H&E reconstruction: H&E→z→H&E; (2) H&E-to-CyCIF translation: H&E→z→CyCIF; (3) CyCIF reconstruction: CyCIF→z→CyCIF; and (4) CyCIF-to-H&E translation: CyCIF→z→H&E. Models were trained with a batch size of 16 and a learning rate of 0.0001 for 100 epochs. Best models were selected based on the lowest validation loss at each epoch end and were then used for downstream application to held-out H&E WSIs. We also experimented with U-Net-like architecture with skip connections between encoder and decoders but found that loss gradients did not propagate to the most internal layers of these models such that meaningful latent representations were not learned.

**Table 2:**
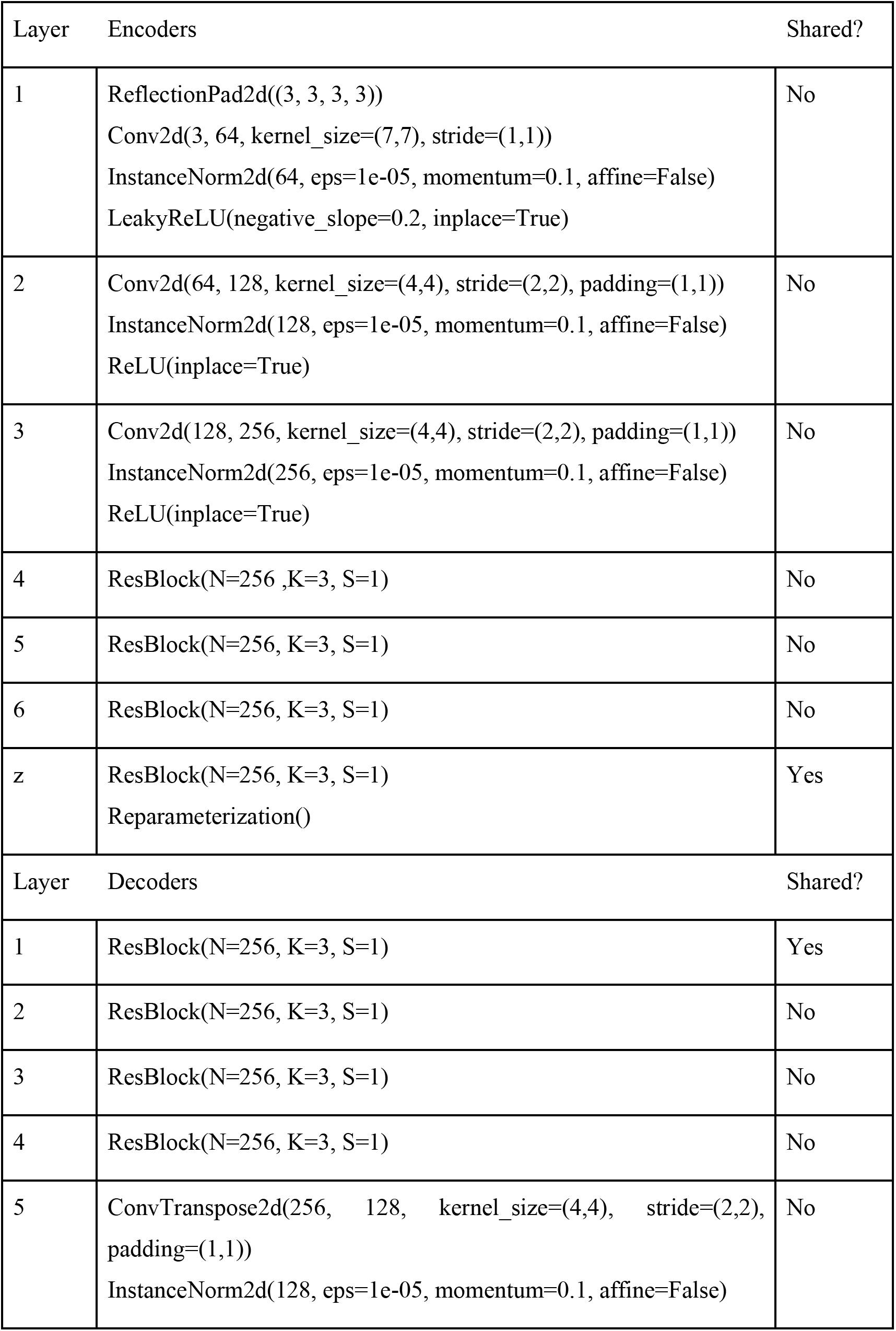
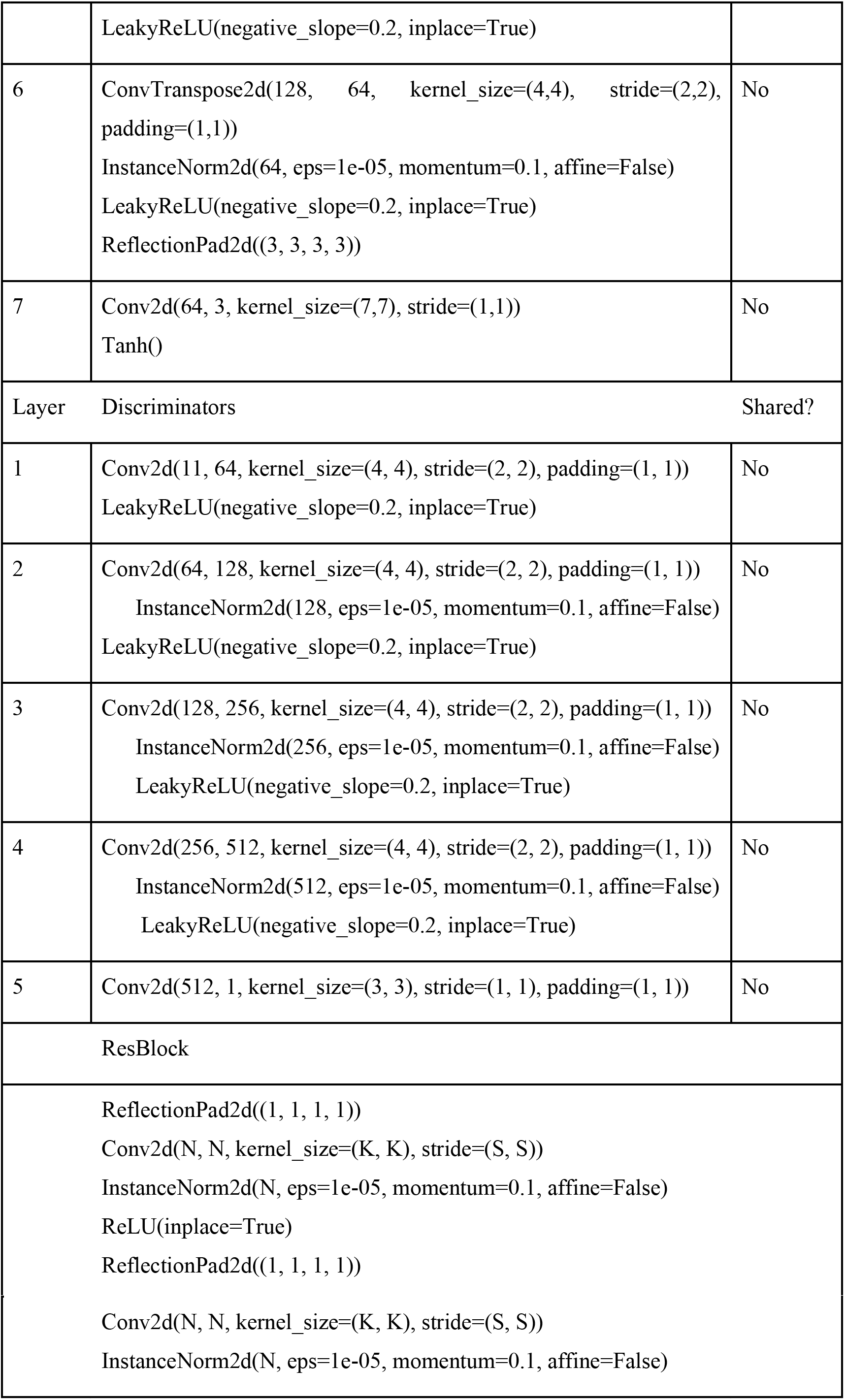
architectures of XAE models.

### Tile cluster identification

Ultimately, we want to evaluate whether deep learning architectures can recapitulate the biological information of both cell type and pathologist, but since VAEs and XAEs operate on a tile by tiles basis, it is necessary to cluster tiles based on their cell type composition. For every tile in the WSI, a vector was created that represented the composition of cell types. The ground truth cell type information was made by K-means clustering these composition vectors (Figure 3). Using the elbow method, we determined that 7 clusters were optimum for evaluation. A smaller number of clusters within the elbow was chosen to better match the number of pathologist annotations for consistency in evaluation. Pathologist information was created manually by an expert pathologist, resulting in 6 distinct tissue types (Figure 3). Tiles were assigned a ground truth tissue type based on the maximum pixel-wise tissue type within the region. 7 clusters were computed for both the standard VAE and the XAE encoding vectors to evaluate against the cell type ground truth clusters.

Several metrics were used to evaluate the ground truth recapitulation. Cluster purity was used to evaluate how well the two methodologies were able to reconstruct the same clusters as ground truth:

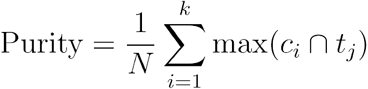

where *N* is the number of data points, *k* is the number of clusters, *c_i_* is the set of predicted clusters and *t_j_* is the set of ground truth clusters. The sklearn implementation of Normalized Mutual Information (NMI)^29^ was used as another metric to evaluate the same question:

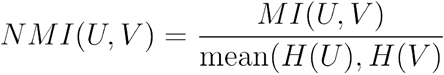

where *U* and *V* are the predicted and ground truth cluster labels, and *H*(*U*) and *H*(*V*) represent the entropy of *U* and *V* respectively. The predicted tile-type clusters were paired to ground truth cell-type clusters and annotations using the Spearman correlation.

To evaluate whether the deep learning models capture the same level of feature information as CyCIF staining, we used the pyrcca^37^ implementations of canonical correlation on the encoded latent feature space and the paired CyCIF tile-wise expressions. The outputs from this process produced two components shared between the two modalities. Quantitatively the correspondence of the two modalities can be measured by the canonical correlation of each component, and qualitatively the correspondence can be observed by the overlap in the scatter plot of the new components.

### Region of Interest (ROIs) selection

#### A. Random Sampling

Random sampling was conducted by randomly drawing a new non-overlapping ROI repeatedly. For bulk analysis and comparison, 1000 random combinations of *k* ROIs were selected where *k* is the number of ROIs found to be optimal for the other sampling methods.

#### B. Convex optimization on composition

If *b* represents counts of cells across the clustered group and *a_i_* represents the cell number belonging to the *i*-th ROI, by solving

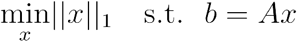

we could identify the minimum number of ROIs to match the WSI cellular population (the main issue of this approach is that we often select homogenous cell populations) where *b* ∈ ℝ^*N*^, *b* represents composition vector of clustered groups within the WSI, each column of *A* ∈ ℝ^*N*×*M*^ represents a possible ROI and each row contains the percentage of tiles in that ROI for each cluster; *N,M* represent the number of clusters and the number of possible ROIs in the WSI respectively.

Since we do not have cell composition beforehand, we will use cluster results based on the latent representation of tiles within ROIs via embedding both H&E and CyCIF. The underlying assumption here is that H&E/CyCIF embedding reflects tile-based cell composition as shown in Figure 2. For the optimization of cluster composition, we solve the optimization problem:

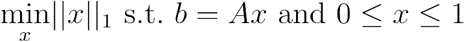

Implementation of this function was conducted using the intlinprog function in MATLAB. The threshold of 0.01 was applied to *x* as a cutoff for selecting relevant ROIs to guarantee all selected ROIs made a significant contribution.

#### C. Convex Optimization with Entropy

To optimize both composition and ROI heterogeneity, we take the entropy of the composition vector into account using the convex optimization function:

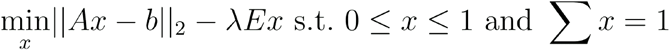

where *E* ∈ ℝ^*M*^ represents the vector of entropies and λ is a hyperparameter governing the weight of entropy. In this experiment, λ was set to 1. Implementation of this function was conducted using CVX^38^ in MATLAB. *x* as the cutoff for selecting relevant ROIs to guarantee all selected ROIs made a significant contribution. The threshold of 0.01 was applied to *x* as a cutoff for selecting relevant ROIs to guarantee all selected ROIs made a significant contribution.

#### D. Evaluation

The quality of the selected representative ROIs was evaluated based on three metrics: Mean squared error (MSE) compared to WSI composition; Jensen-Shannon Divergence (JSD) of the ROI and WSI compositions; and mean ROI entropy. Mean squared error was calculated using:

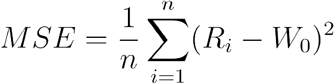

where *n* is the number of predicted clusters, *R* is the percent composition of each cluster within all selected ROIs combined, and *W*_0_ is the percent composition of each cluster within the WSI. JSD was calculated using:

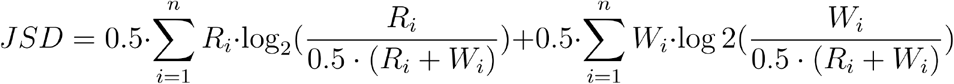

where *n* is the number of predicted clusters, *R* is the percent composition of each cluster within all selected ROIs combined, and *W* is the percent composition of each cluster within the WSI. The mean entropy was calculated using:

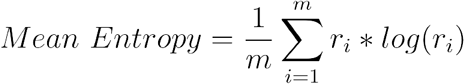

where *m* is the number of selected ROIs and *r_i_* is the percent composition within each individual ROI.

## ACKNOWLEDGEMENTS

This work was carried out with major support from National Cancer Institute (NCI), Human Tumor Atlas Network (HTAN) Research Centers at OHSU (U2CCA233280), and Harvard Medical School (U2CCA233262). Y.H.C is supported by R01 CA253860 and Kuni Foundation Imagination Grants. The resources of the Exacloud high-performance computing environment developed jointly by OHSU and Intel and the technical support of the OHSU Advanced Computing Center is gratefully acknowledged.

## COMPETING INTERESTS

J.W. Gray has licensed technologies to Abbott Diagnostics; has ownership positions in Convergent Genomics, Health Technology Innovations, Zorro Bio, and PDX Pharmaceuticals; serves as a paid consultant to New Leaf Ventures; has received research support from Thermo Fisher Scientific (formerly FEI), Zeiss, Miltenyi Biotech, Quantitative Imaging, Health Technology Innovations, and Micron Technologies.

P.K. Sorger is a member of the SAB or BOD member of Applied Biomath, RareCyte Inc. and Glencoe Software, which distributes a commercial version of the OMERO database; P.K. Sorger is also a member of the NanoString SAB. In the last 5 years, the Sorger laboratory has received research funding from Novartis and Merck.

The other authors declare no outside interests.

## DATA and SOFTWARE AVAILABILITY

All full-resolution images derived image data (e.g., segmentation masks) and all cell count tables will be publicly released via the NCI-sponsored repository for Human Tumor Atlas Network (HTAN; https://humantumoratlas.org/) at Sage Synapse.

All software used in this manuscript is detailed in the article’s Methods section and its Supplementary Information. The associated scripts are freely available via GitHub as described at https://gitlab.com/eburling/SHIFT.

## SUPPLEMENTARY FIGURES

**Supp. Figure 1:**
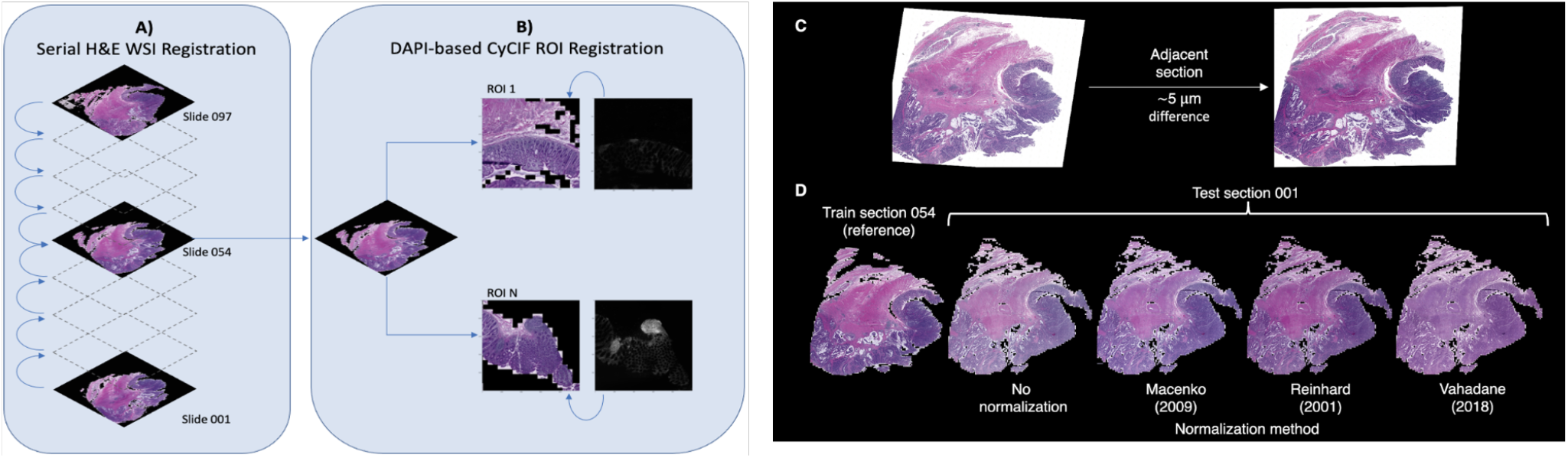
Registration schema for 3D H&E and CyCIF imaging dataset and normalization overview. (A) To register H&E in the three-dimensional setting, we sequentially registered all slides to the center using the transforms propagated from previous layers. (B) CyCIF was then finely registered to the adjacent H&E images at the ROI level to maximize single-cell level correspondence. Registration of CyCIF and H&E was performed using binarized DAPI and thresholded H&E to align nuclei. (C) Tissue sections are subject to technical variability in stain intensity, even between adjacent sections that are separated by only ~5 μm. (D) Representative results of H&E stain normalization. The stain intensity distribution of the test section 001 is transformed to match that of the reference section 054 which was used for SHIFT model training.

**Supp. Figure 2:**
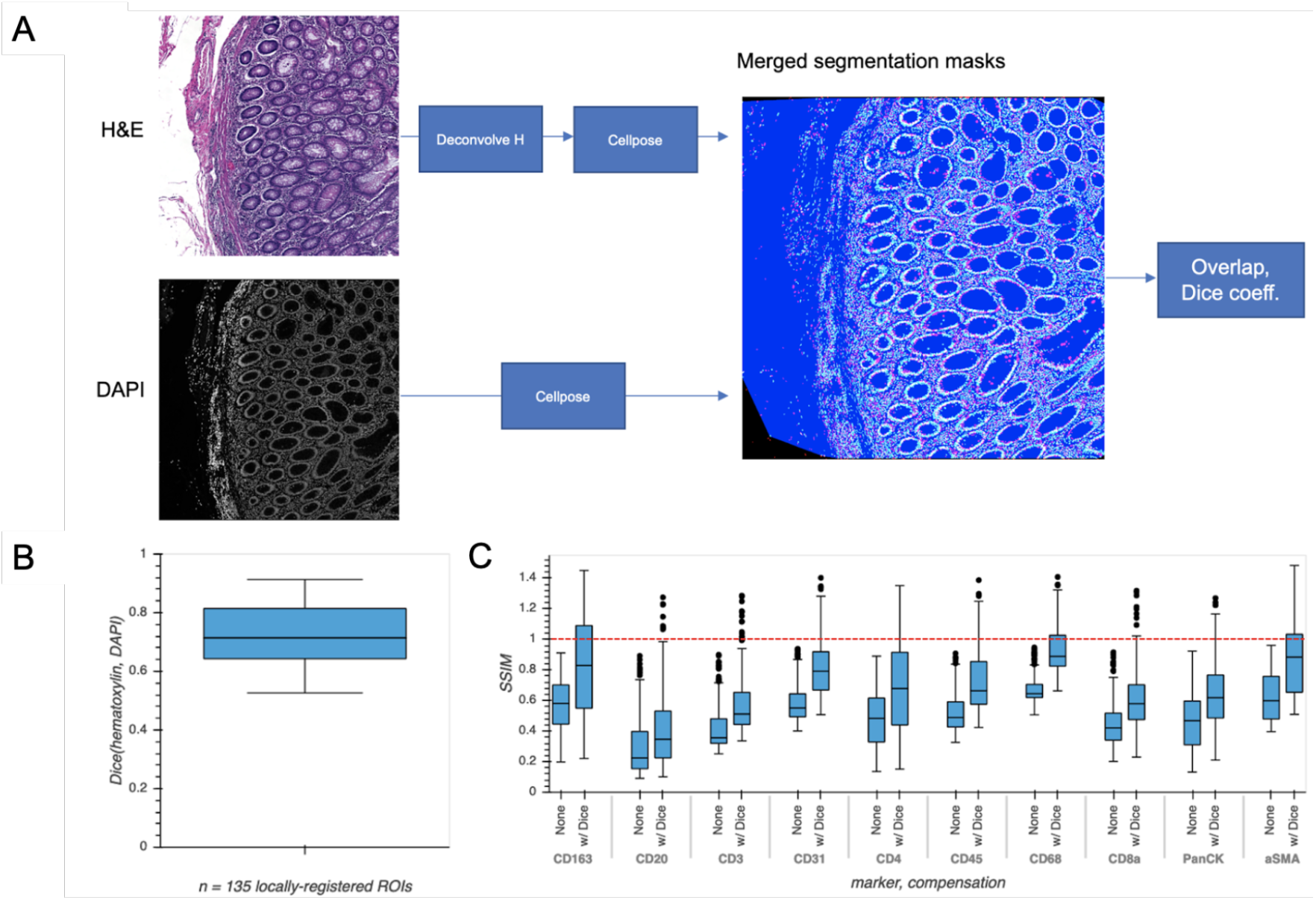
Estimating upper bound on SHIFT performance by measuring concordance between nuclei overlap in adjacent sections for locally-registered ROIs from H&E/CyCIF test sections 096/097. (A) The Dice coefficients describing the overlap of nuclear masks from ROIs of adjacent sections were used as compensation factors for evaluating virtual stains. (B) Boxplot describing the distribution of Dice coefficients of the 135 locally-registered ROIs from H&E/CyCIF test sections 096/097. (C) Boxplots describing the distributions of structural similarity (SSIM) of real vs. virtual CyCIF ROIs over the 135 locally-registered ROIs from H&E/CyCIF test sections. The red dotted line indicates the unity line describing Dice-compensated SSIM.

**Supp. Figure 3:**
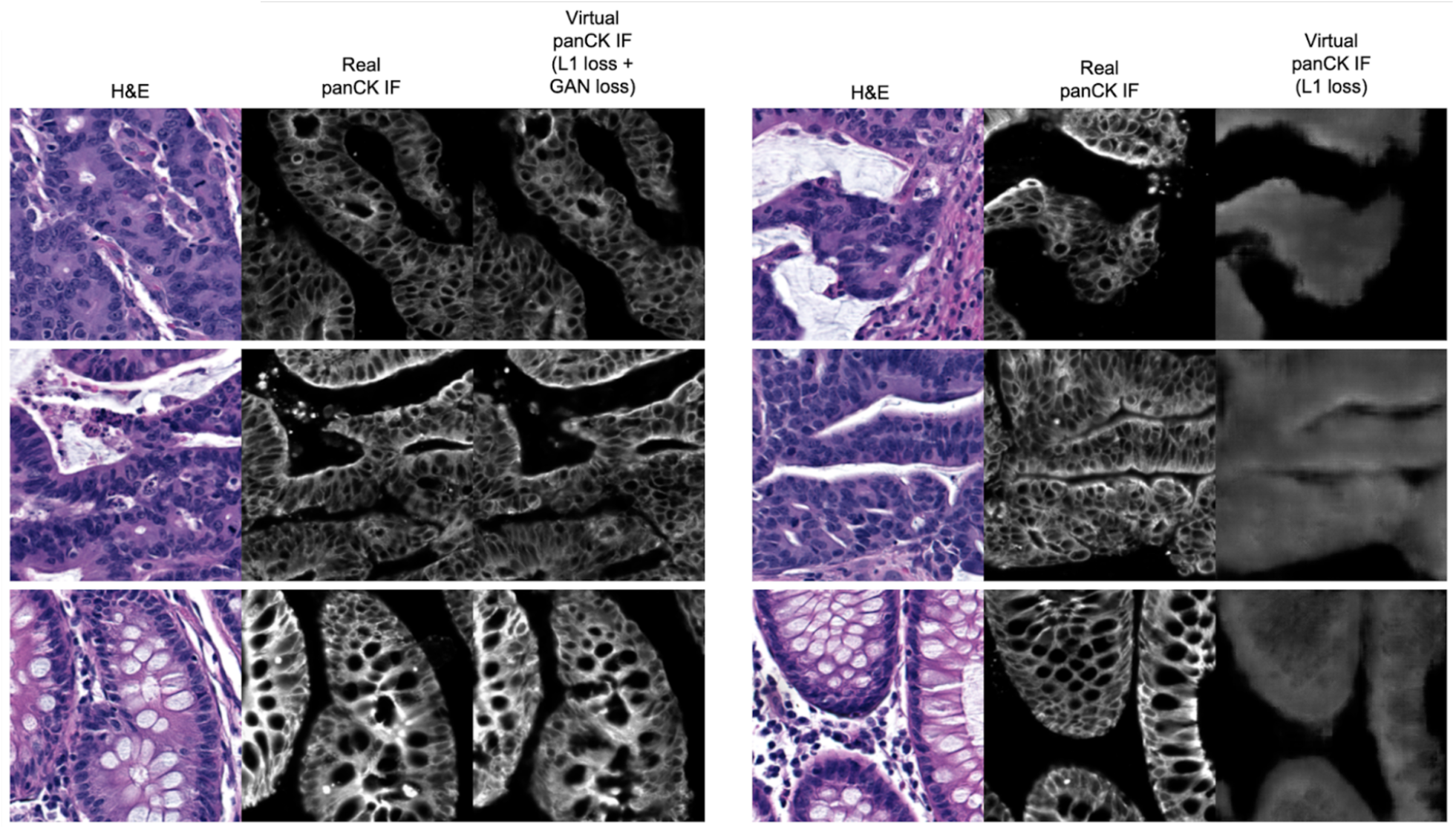
Virtual staining outcomes with different loss functions. The left panels correspond to results from the full SHIFT model (generator and discriminator) and the right panels correspond to results from a model consisting of a generator only.

**Supp. Figure 4:**
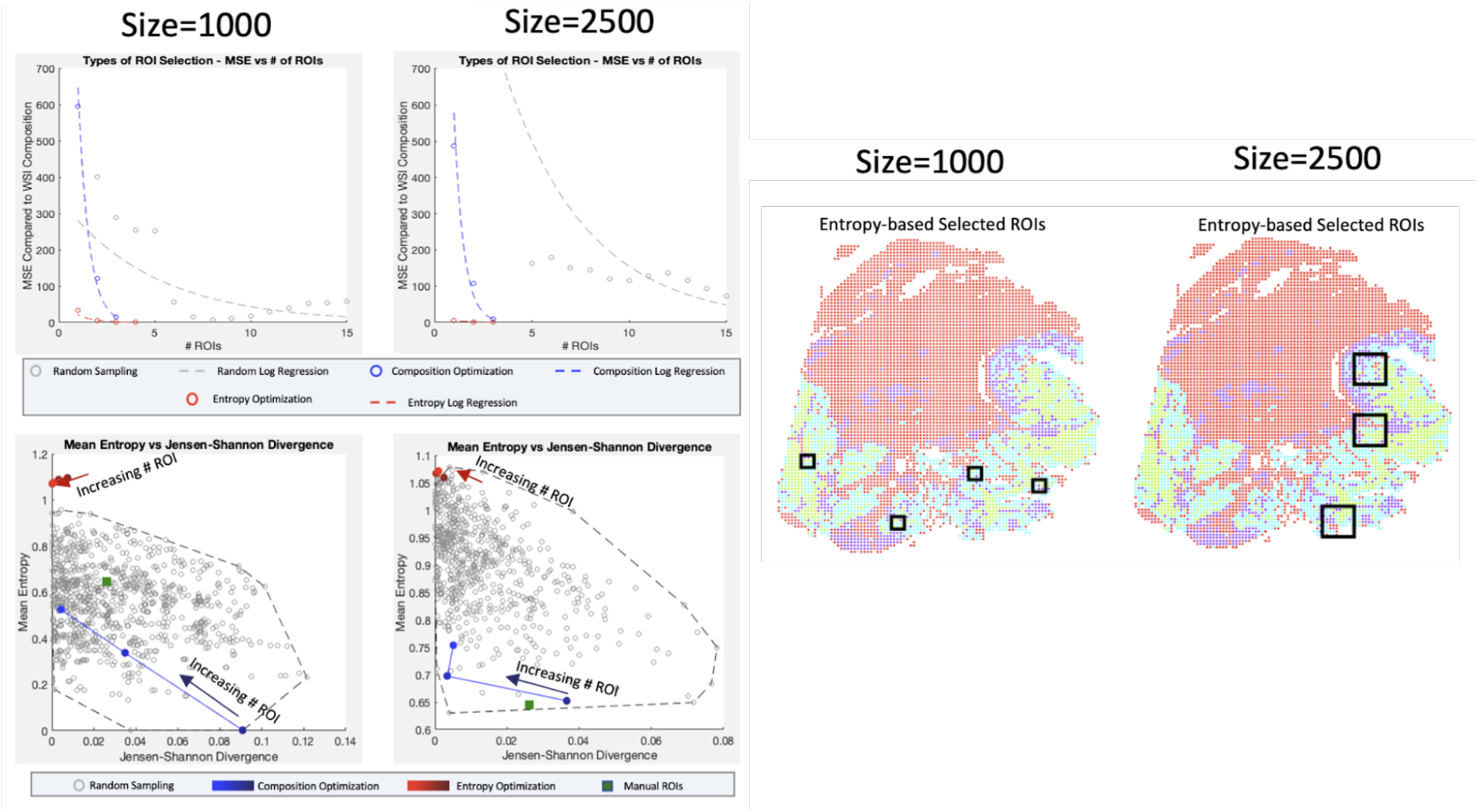
Optimization of ROI selection within a restricted set of cells. For two ROI sizes (1000 pixel, 2500 pixel) and three sampling techniques (random sampling, convex optimization using cell type composition, convex optimization using cell type composition, and regional entropy), we calculate the optimal selection of ROI. Before calculation, we restrict the cell types of interest to only tumor and immune type cells targeted by manual annotation. **(Top row)** By calculating the MSE for a range of ROI, we can evaluate each technique’s rate and quality of convergence. (**Middle row)** Selections of representative ROIs are evaluated based on two metrics (Entropy for tissue heterogeneity and Jensen-Shannon Divergence for composition similarity.) Random sets of 7 ROIs are generated 1000 times to portray the baseline pattern. Selections from convex optimizations are plotted with increasing numbers of ROIs to show the change in performance. The performance of the manually selected ROIs is also shown to emphasize the bias in targeted sampling. **(Bottom row)** The optimal ROIs are shown for convex entropy optimization at each size of ROI. Image colors portray the XAE-labeled cell types (red being cell types not considered in this analysis).

